# Tertiary lymphoid structure-related immune infiltrates in NSCLC tumor lesions correlate with low tumor-reactivity of TIL products

**DOI:** 10.1101/2024.02.19.580998

**Authors:** Suzanne M. Castenmiller, Nandhini Kanagasabesan, Aurélie Guislain, Benoît P. Nicolet, Marleen M. van Loenen, Kim Monkhorst, Alexander A.F.A. Veenhof, Egbert F. Smit, Koen J. Hartemink, John B.A.G. Haanen, Rosa de Groot, Monika C. Wolkers

## Abstract

Adoptive transfer of tumor infiltrating lymphocytes (TIL therapy) has shown great potential for the treatment of solid cancers, including non-small cell lung cancer (NSCLC). However, not all patients benefit from this therapy, and the parameters that define the likelihood of TIL products to be tumor reactive are to date unknown. Defining prognostic markers that correlate with high level of tumor-reactivity is key for achieving better tailored immunotherapies.

To determine whether the composition of immune cell infiltrates correlates with the tumor-reactivity of expanded TIL products, we employed multi-parameter flow cytometry to characterize the immune cell infiltrates from 26 early-stage, and 20 late-stage NSCLC tumor lesions. Unbiased flow cytometry analysis with Cytotree and Spearman’s Rank Correlation was used to correlate immune infiltrates with the expansion rate, immune cell activation and T cell differentiation state, and the anti-tumor response of TIL products generated from the same lesions.

The composition of tumor immune infiltrates was highly variable between patients, irrespective of the disease stage. High percentages of B cell infiltrates positively correlated with the presence of conventional CD4^+^ T cells, and an overall increase of naïve T cell infiltrates. In contrast, high B cell infiltrates negatively correlated with the tumor-reactivity of expanded TIL products, as defined by cytokine production upon exposure to autologous tumor digest. Tumors with high B cell infiltrates contained IgD^+^BCL6^+^ B cells and CXCR5^+^BLC6^+^ CD4^+^ T cell infiltrates and an increased percentage of naïve CD8^+^ T cells, indicative of the presence of tertiary lymphoid structures (TLS) in tumors with high B cell infiltrates.

This study reveals that the composition of immune cell infiltrates in NSCLC tumors associates with the functionality of expanded TIL products from NSCLC tumor lesions. Importantly, the tumor-responsiveness of TIL products negatively correlated with the presence of TLS-associated immune infiltrates in tumors. Our finding may thus help improve patient selection for TIL therapy.

## Introduction

Adoptive cellular therapy with tumor infiltrating lymphocytes (TILs) has shown high potential for the treatment of solid tumors^1^. The exciting findings from small-scale phase I trials for advanced melanoma patients receiving TIL therapy^2–6^ were recently confirmed by a larger scale phase III clinical trial^7^, revealing the great treatment potential of TIL therapy for metastasized melanoma patients. Importantly, and as previously suggested by a small-scale study^8^, this multi-center open-label trial demonstrated that not only treatment-naïve patients responded to TIL therapy. It also showed efficacy in patients that were resistant to checkpoint inhibitors^7^.

Owing to these successes, the generation of tumor-reactive TIL products has been assessed for other solid tumors, including bladder cancer^9^, ovarian cancer^10^, and renal cell cancer^11^. Additionally, we and others showed that TIL products generated from treatment-naïve non-small cell lung cancer (NSCLC)-patients contain poly-functional tumor-reactive T cell products^12,13^. As for melanoma alike^7,8^, we found that tumor-reactive T cells were also present in TIL products from late-stage pretreated NSCLC patients, indicating that TIL therapy could potentially be broadly applicable to NSCLC patients^14^. In line with these findings, a first very promising phase I clinical trial with TIL therapy in PD1-refractory patients recently reported a reduced tumor burden in most patients^15^. Consequentially, more clinical trials with TIL therapy are expected to start in the near future^16,17^.

However, similar to treatments with checkpoint inhibitors^18–21^, not all patients respond equally well to TIL therapy. For metastatic melanoma, 10-20% of the patients undergo long-term complete remission^6,10^. Others, however, undergo only partial remission or stable disease, or do not benefit from TIL therapy at all^4,6,7,15,22^. To date, reports on indicators that stratify patients with a high likelihood to generate tumor-reactive TIL products are scarce. Such prediction tools are thus in high demand, as they will help select the patients who will benefit most from TIL therapy.

The divergent response rate to treatment with checkpoint inhibitors has been attributed to the immune cell composition of the tumor microenvironment (TME)^21,23,24^. In particular high percentages of B cell infiltrates, and the presence of tertiary lymphoid structures (TLS) have been indicated as positive indicators of responding to checkpoint inhibitors^23,25^. Also for TIL therapy, first indications of the influence of the TME composition in generating TIL products have been collected. In melanoma patients, the T cell differentiation status was proposed to correlate with the efficacy of the TIL product^5,26^, and in a small cohort, a correlation with the activation status of macrophages and dendritic cells was observed^27^. For NSCLC-derived TIL products, such correlations with the immune cell infiltrates in tumor lesions have, to our knowledge, not yet been reported. We hypothesized that studying the immune cell compartment in NSCLC tumor digests at baseline could potentially provide important insights into the quality and tumor-reactivity of expanded TIL products.

In this study, we mapped the immune cell infiltrates from 26 treatment-naïve early-stage and from 20 pre-treated late-stage NSCLC patients. Even though tumor-specific immune infiltrates were enriched for lymphoid cells, we observed a high inter-patient variation. In particular B cells and conventional CD4^+^ T cells co-inhabited tumor lesions. Surprisingly, whereas NKT cells and neutrophils displayed a positive correlation with the presence of tumor-reactive T cells in expanded TIL products, high percentages of B cells and conventional CD4^+^ T cells negatively correlated with the functionality of the TIL products. Intriguingly, tumors with a high B cell infiltrates contained BCL6^+^ mantle zone B cells and CXCR5^+^ follicular helper T cells (Tfh), strongly indicating the presence of tertiary lymphoid structures (TLS). We therefore postulate that the presence of TLS-related immune infiltrates negatively impacts the tumor-reactivity potential of expanded TIL products.

## Results

### Tumor lesions display an influx of lymphoid cells, irrespective of the disease stage

To study the composition of immune cell infiltrates in NSCLC tumor lesions, we collected tumor and adjacent lung tissue from 26 early-stage, treatment-naïve patients, and metastatic tumor lesions from 20 late-stage NSCLC patients. The late-stage patients had different histories of treatment, including chemotherapy, immunotherapy, chemo-immuno combination therapy, or small-molecule inhibitor therapy (EGFR and ALK inhibitors; table 1). 27 female and 19 male patients between 37 and 87 years of age (mean 63.2 years) were included, of which 37 patients (78%) had a history of smoking. Tumor size ranged from 0.7-8 cm^2^, with an average of 3.5 cm^2^ (table 1).

We obtained more viable cells per gram tissue from enzymatically digested early-stage tumor lesions (32.3×10^6^ ± 23.8×10^6^) than from the paired adjacent lung tissue (16.0×10^6^ ± 25.3×10^6^, p=0.0013; table 1, figure 1A). Tissue digests from late-stage tumors contained comparable numbers of viable cells per gram tissue cell to early-stage tumors (34.1×10^6^ ± 34.6×10^6^; table 1, figure 1A). The lung digest contained higher percentages of CD45^+^ immune cell infiltrates (29.6% ± 18.5%) than both early- and late-stage tumors (21.4% ± 20.0% and 16.0% ± 20.1%, respectively; p=0.024; figure 1B). Nevertheless, the number of CD45^+^ cells isolated per gram tissue was similar from lung and tumor digests (figure 1C).

**Figure 1.**
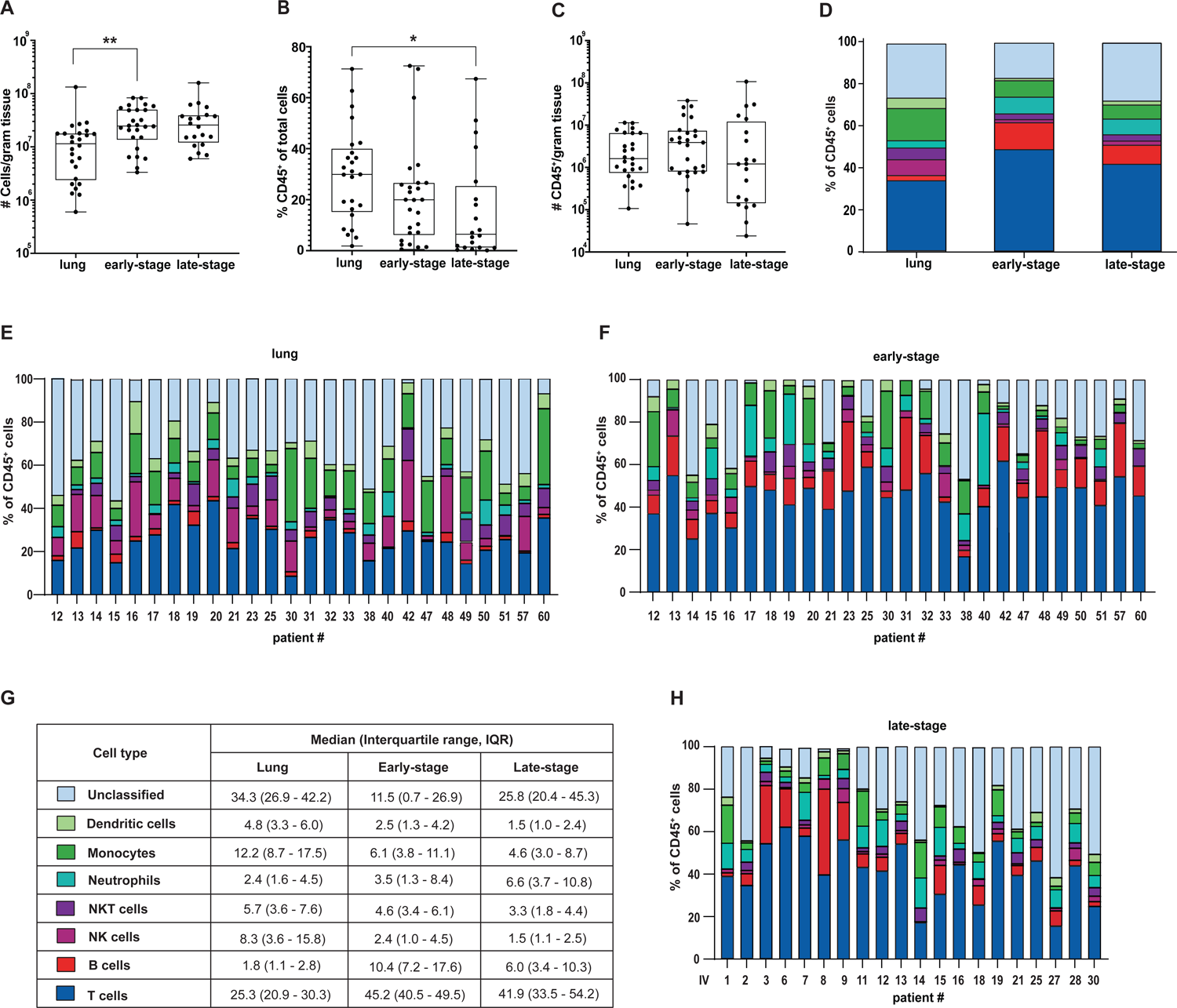
Increased T and B cell infiltrates in NSCLC tumor lesions with high inter-patient variability. (A) Cells per gram tissue obtained from lung tissue (left; n=26), early-stage NSCLC tumor lesions (middle; n=26) and late-stage NSCLC tumor lesions (right; n=20) after digestion. (B, C) CD45^+^ immune infiltrates as (B) percentage of total number of cells, and (C) per gram tissue from indicated tissue digests. (D) Averaged percentages of various immune infiltrates of indicated tissue digests, immune cell types are colored according to cell type shown in panel G. (E-H) Composition of immune infiltrates in (E) lung tissue, (F) early-stage and (H) late-stage NSCLC tumor lesions from individual patients. (G) Percentages of immune infiltrates from E-H are presented as median with interquartile range. Statistical significance for lung and early-stage tumor digest was defined with paired t-test and for late-stage tumors compared to early-stage tumor and lung tissue with unpaired t-test. p-value *>0.05, **>0.005.

To define the percentage of lymphoid (T cell, B cell, NK cell, NKT cell) and myeloid (neutrophil, monocyte, dendritic cell) infiltrates, we performed an integrated multi-parameter flow cytometry analysis followed by a multistep analysis in R as described in the methods (table 2, supplemental figure 1, supplemental figure 2A). CD45^+^ immune cells that could not be attributed to one specific cell subset based on the antibody panel we used (table 2) were defined as ‘unclassified’. The percentage of unclassified CD45^+^ cells (which should include e.g. basophils, mast cells, eosinophils and possibly MAIT cells) was substantially reduced in early-stage tumor lesions compared to lung tissue, a feature that was not sustained in late-stage tumors (figure 1D,G). Overall, early- and late-stage tumor lesions were more alike than early-stage tumors with the paired lung tissue (figure 1D). In accordance with previous studies^41,42^, we found an increase of T cells and B cells in tumor lesions, at the cost of lower percentages of myeloid and NK cell infiltrates in the tumor lesions (figure 1D-H).

Importantly, we observed a high interpatient variability in immune infiltrate composition (figure 1E-H). For example, the interquartile range of B cell infiltrates in late-stage tumors ranged from 3.4% to 10.3% (figure 1G), but it could be as little as 0.3% and as much as 44.6% in patients. Similarly, neutrophil infiltration ranged from 0.1% to 33.8%. Thus, tumor lesions display increased T and B cell infiltrates compared to healthy lung, yet the overall composition of immune infiltrates in NSCLC tumor lesions is highly variable between patients, irrespective of their disease stage.

### B cell infiltration positively correlates with CD4^+^ conventional T cell infiltrates in tumor lesions

Because immune cell infiltrates communicate with each other^24^, we examined how the presence of one immune cell subset related with that of other subsets, using Spearman’s correlation coefficient (r_s_). We first examined the lung and tumor lesions individually (supplemental figure 2). For lung tissue, we specifically found that CD8**^+^** T cell infiltration in lung tissue positively correlated with the presence of Tregs and DCs (supplemental figure 2B,C). Also, in early-stage and in late-stage NSCLC tumor lesions, we found a number of correlations between immune cell infiltrates (supplemental figure 2D-G). To identify the correlations of immune infiltrates that were common between early- and late-stage tumors, we re-analyzed the relation between immune infiltrates with the two NSCLC cohorts combined. The positive correlation between neutrophil and monocyte infiltrates (r_s_=0.35, p=0.017) and between B cell and conventional CD4^+^ T cell infiltrates (r_s_=0.43, p=0.0042) was conserved in this combined analysis (figure 2A,B; supplemental figure 2H), suggesting a tumor-intrinsic correlation between different NSCLC-infiltrating immune cell subsets.

**Figure 2.**
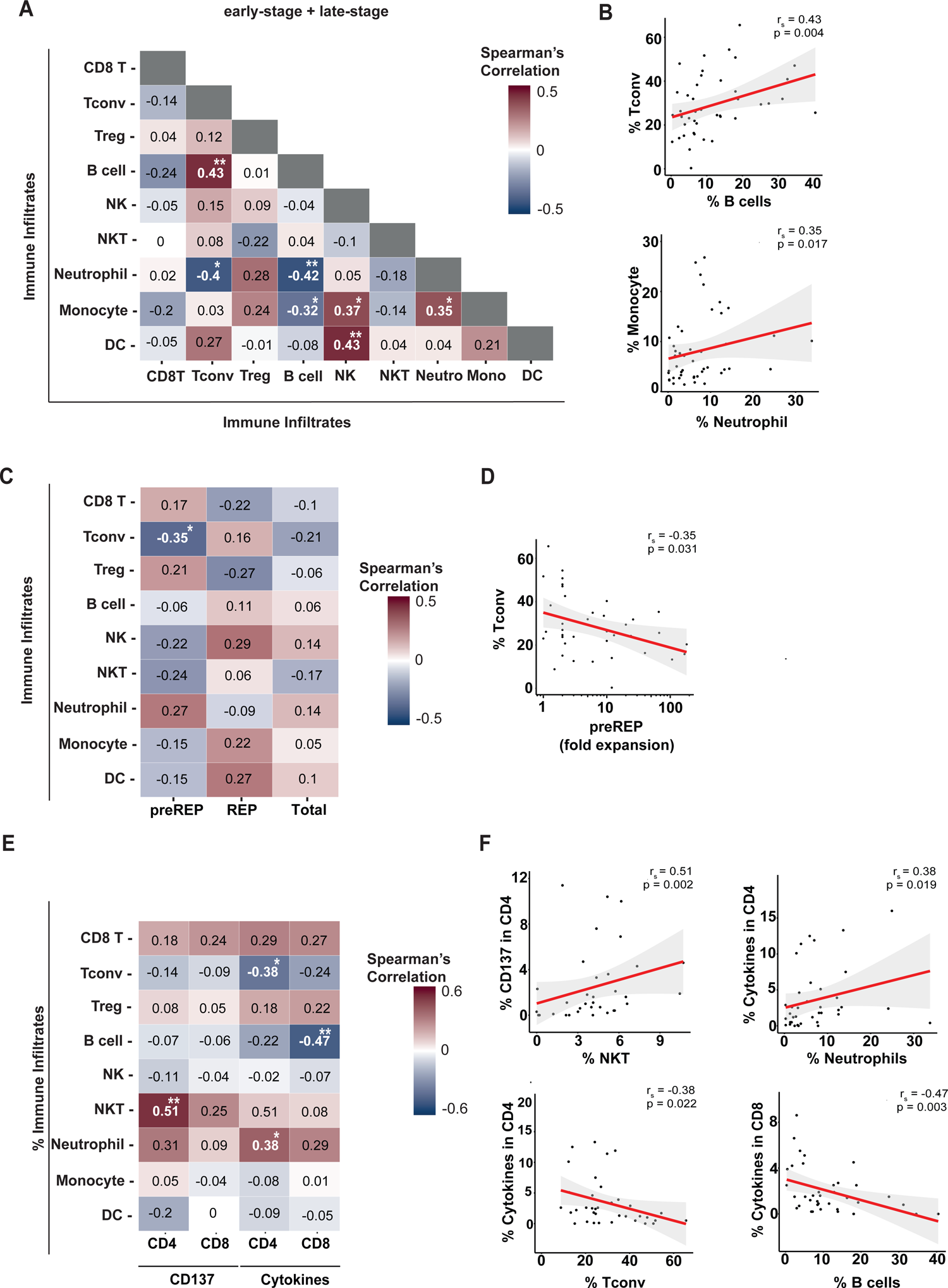
Tumor-specific immune infiltrates correlate with T cell functionality of expanded TIL products. (A) Correlation heatmap between different immune infiltrates of early- and late-stage NSCLC tumor lesions combined (n=46). (B) Scatterplots show significant correlations between B cell infiltrate with conventional CD4^+^ T cells, and monocyte infiltrate with neutrophils. (C) Correlation heatmap between the percentage of immune infiltrate subsets of early-stage and late-stage NSCLC tumor lesions combined with the expansion rate (fold change) of T cells during preREP (first 1-13 days), REP (second 1-13 days) and total expansion phase (preREP + REP). (D) The scatterplot shows the correlation between the percentage of conventional CD4^+^ T cells with the preREP phase. (E) Correlation heatmap of indicated immune infiltrates of NSCLC tumor lesions ex vivo with anti-tumoral response of expanded TIL products, as defined by CD137 expression, or production of at least one of the key cytokines (IFNγ, TNF, and/or IL-2) by CD4^+^ and CD8^+^ T cells, after 6-7h of co-culture with autologous tumor digest. (F) Scatterplots show significant correlations from panel (E) depicted for individual patients. All correlation heatmaps report Spearman’s coefficient (r_s_) and *p<0.05, **p<0.005. All scatterplots show the Spearman’s coefficient (r_s_) and p-value. The red line indicates the regression line of the correlation, and the grey area represents the confidence interval.

### B and NKT cell infiltrates in the TME correlate with tumor-reactivity of expanded TIL products

Successful TIL therapy requires sufficient expansion to achieve a clinical response^2^. To define whether the cellular composition of the tumor digest correlated with the capacity of TILs to expand *in vitro*, we used TIL expansion data we previously reported^12,14^, and complemented them with the analysis for tumors from newly recruited patients (table 1). We observed no overt correlations of the percentage of immune infiltrates with the fold expansion rate during the pre-REP (first 10-13 days of expansion) and the REP phase (second 10-13 days of expansion), except from a slight negative correlation of conventional CD4^+^ T cells with TIL expansion during the pre-REP phase (r_s_=-0.35, p=0.031; figure 2C,D). Overall, the composition of immune infiltrates in digested tumor lesions had a limited effect on the TIL expansion from the same lesion.

Another key objective for effective TIL therapy is the presence of tumor-reactive T cells in the expanded TIL product. We therefore questioned whether any immune cell type present in tumor digest correlated with tumor reactivity of the TIL products, as defined by CD137 (4-1BB) expression and the production of at least one the key pro-inflammatory cytokines (IFNγ, TNF and IL-2) by T cells upon exposure to autologous tumor digest for 6-7 hours (table 1). Interestingly, the percentage of NKT cells and neutrophils in the TME positively correlated with the CD137 expression on expanded CD4^+^ TILs (r_s_=0.51, p=0.0016, and r_s_=0.38, p=0.019, respectively; figure 2E,F). Surprisingly, however, the percentage of conventional CD4^+^ T cells in the TME negatively correlated with the cytokine production of expanded CD4^+^ TILs (r_s_= −0.38, p=0.022) (figure 2E,F). Even more prominently, the percentage of B cells in the TME negatively correlated with the cytokine expression of expanded CD8^+^ TILs (r_s_=-0.47, p=0.0031) (figure 2E,F). A similar trend was observed for CD4^+^ TILs (figure 2E). Thus, TIL products that were generated from tumor lesions with high B cell and conventional CD4^+^ T cell infiltrates correlated with TIL products displaying lower percentages of tumor reactivity.

### Presence and activation status of neutrophils negatively correlates with tumor-reactive TIL products

The activation status of TME infiltrates has been reported to influence the presence and functionality of tumor-reactive T cells^43–46^, a feature that was also reported for NSCLC^42,47^. PD-L1, which is expressed on activated immune cells, plays a key role in suppressing inflammatory immune responses^48^. Likewise, HLA-DR is expressed on activated antigen presenting cells such as DCs, monocytes and B cells, and on activated T cells^47,49,50^. We questioned whether the PD-L1 or HLA-DR expression on the immune infiltrates could explain the high variety of anti-tumor responses we observe in TIL products. In NSCLC infiltrates, DCs and monocytes displayed high percentages of PD-L1 and HLA-DR expression, irrespective of the disease stage (supplemental figure 3A, figure 3A,B). B cells primarily showed high percentages of HLA-DR expression, again in both early- and late-stage tumors (figure 3A,B). For all other cell types, the percentage of PD-L1 and HLA-DR expressing cells was more variable. Of note, PD-L1 or HLA-DR expression on CD3^+^ T cell infiltrates did not correlate with increased Treg infiltration (supplemental figure 3B), indicating that PD-L1 or HLA-DR expression rather originated from conventional CD3^+^ T cells. The most prominent variation of PD-L1 and HLA-DR expression was observed for neutrophils (figure 3A, B). Independently of the tumor stage, neutrophils showed very high, or very low expression of these two activation markers (figure 3A,B,C). As for monocytes and DCs alike, PD-L1 or HLA-DR are mostly co-expressed (figure 3C).

**Figure 3.**
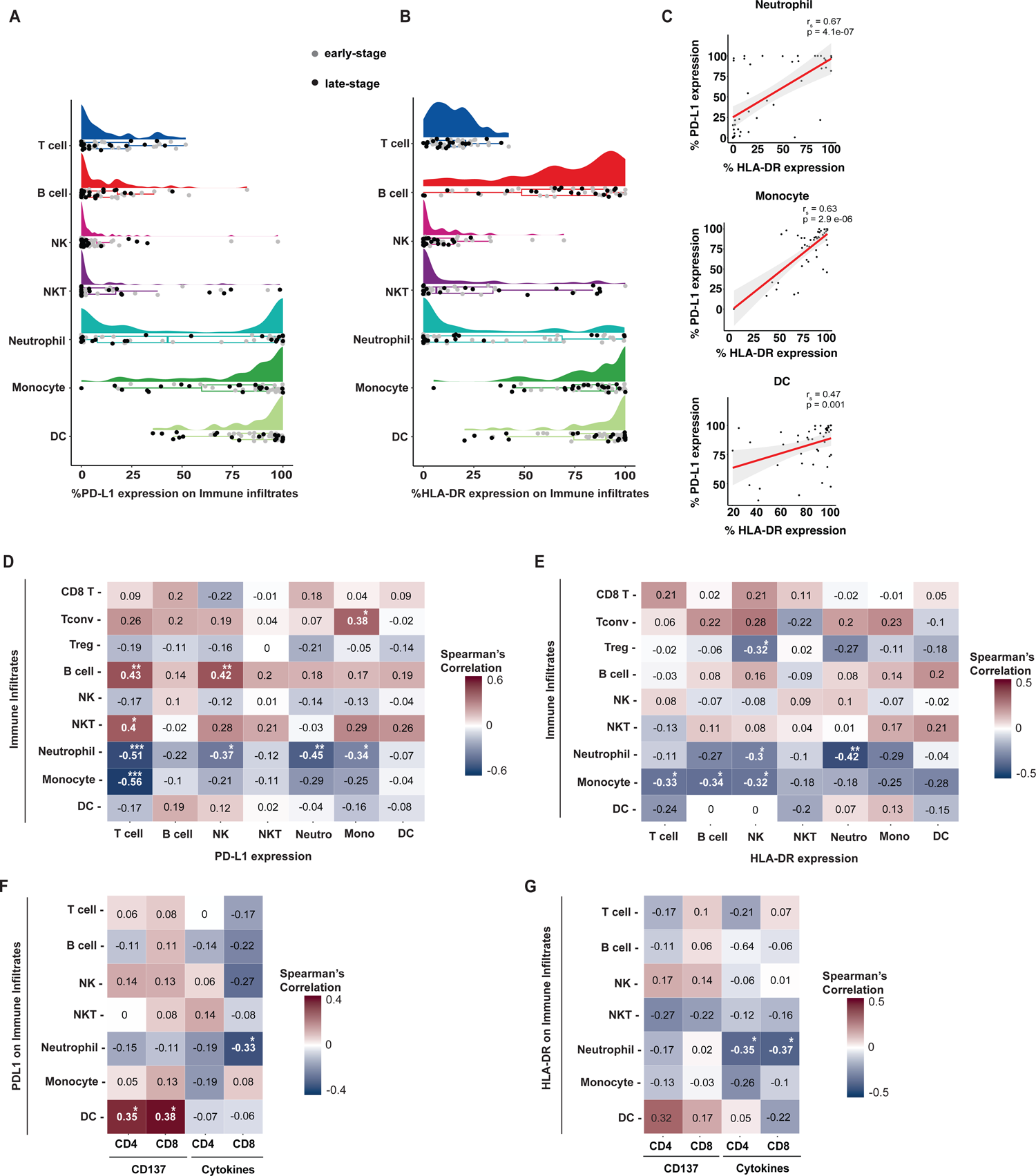
PD-L1 and HLA-DR expression on DCs and neutrophils correlate with T cell functionality. (A, B) Raincloud plots of (A) PD-L1 and (B) HLA-DR expression on different immune cell subsets of early-stage (gray dots) and late-stage (black dots) NSCLC tumors. Per cell type, a density plot (top) and boxplot with each dot corresponding to individual patients is shown. (C) Scatterplots show correlation between PD-L1 and HLA-DR expression on indicated cell types. (D, E) Correlation heatmaps of different immune cell infiltrates with the percentage of (D) PD-L1 expression and (E) HLA-DR expression on different immune infiltrates. (F, G) Correlation heatmaps of different immune cell infiltrates with the percentage of (F) PD-L1 expression and (G) HLA-DR expression with indicated anti-tumoral response of expanded TIL products upon 6h of co-culture with autologous tumor digest. All correlation heatmaps report Spearman’s coefficient (r_s_) and *p<0.05, **p<0.005, ***p<0.0005. All scatterplots show the Spearman’s coefficient (r_s_) and p-value. The red line indicates the regression line of the correlation, and the grey area represents the confidence interval.

We next determined if PD-L1 or HLA-DR expression on immune cells corresponded with the composition of TME infiltrates. B cell infiltrates positively correlated with PD-L1 expression on NKT and T cell infiltrates (figure 3D, supplemental figure 3C). In contrast, neutrophil and monocyte infiltrates negatively correlated with PD-L1 expression on T cells, and neutrophils with the PD-L1 expression on neutrophils, NK cells and monocytes (figure 3D, supplemental figure 3C). The HLA-DR expression correlates with TME infiltrates were less pronounced, yet, neutrophil infiltrates negatively correlated with HLA-DR expression on NK cells and neutrophils (figure 3E, supplemental figure 3D). Thus, the presence of neutrophils in the TME negatively correlates with the immune activation status of other TME immune infiltrates.

We also studied whether the PD-L1 or HLA-DR expression on TME infiltrates correlated with the tumor-reactivity of expanded TILs. We found no effect of PD-L1 and HLA-DR expression on B or T cells with tumor reactivity (figure 3F,G). In contrast, PD-L1-epxressing DCs in the TME positively correlated with CD137 expression on expanded CD4^+^ and CD8^+^ TILs (figure 3F, supplemental figure 3E), suggesting that the presence of activated DCs indicates the presence of tumor-reactive T cells. Interestingly, PD-L1 and HLA-DR expression on neutrophils negatively correlated with cytokine expression of tumor-exposed expanded TILs (figure 3F,G; supplemental figure 3E,F). Thus, not only the overall presence of neutrophils in NSCLC tumors negatively correlates with the percentage of tumor-reactive T cells in TIL products, but also their PD-L1 and HLA-DR expression levels.

### B cells positively correlate with naïve CD8^+^ in the TME, at the cost of effector memory CD4^+^ T cells

The T cell differentiation status was proposed to correlate with *in vivo* efficacy of TIL therapy in melanoma patients^5,26^. Also in NSCLC patients, the T cell differentiation status can greatly vary^51^. To determine how the T cell differentiation status correlated with the tumor-reactivity of expanded TILs in our cohort, we measured the percentage of naïve T cells (Tn), central memory T cell (Tcm), effector memory cells (Tem), and terminally differentiated effector T cells (Temra) *ex vivo*, based on CD27 and CD45RA expression (supplemental figure 4A; table 1). We found a negative correlation of naive CD4^+^ and CD8^+^ T cells with CD137 expression and with cytokine expression of expanded TIL products (figure 4A, supplemental figure 4B), suggesting that high levels of naïve T cells in the TME result in inferior tumor-reactivity of the TIL products.

**Figure 4.**
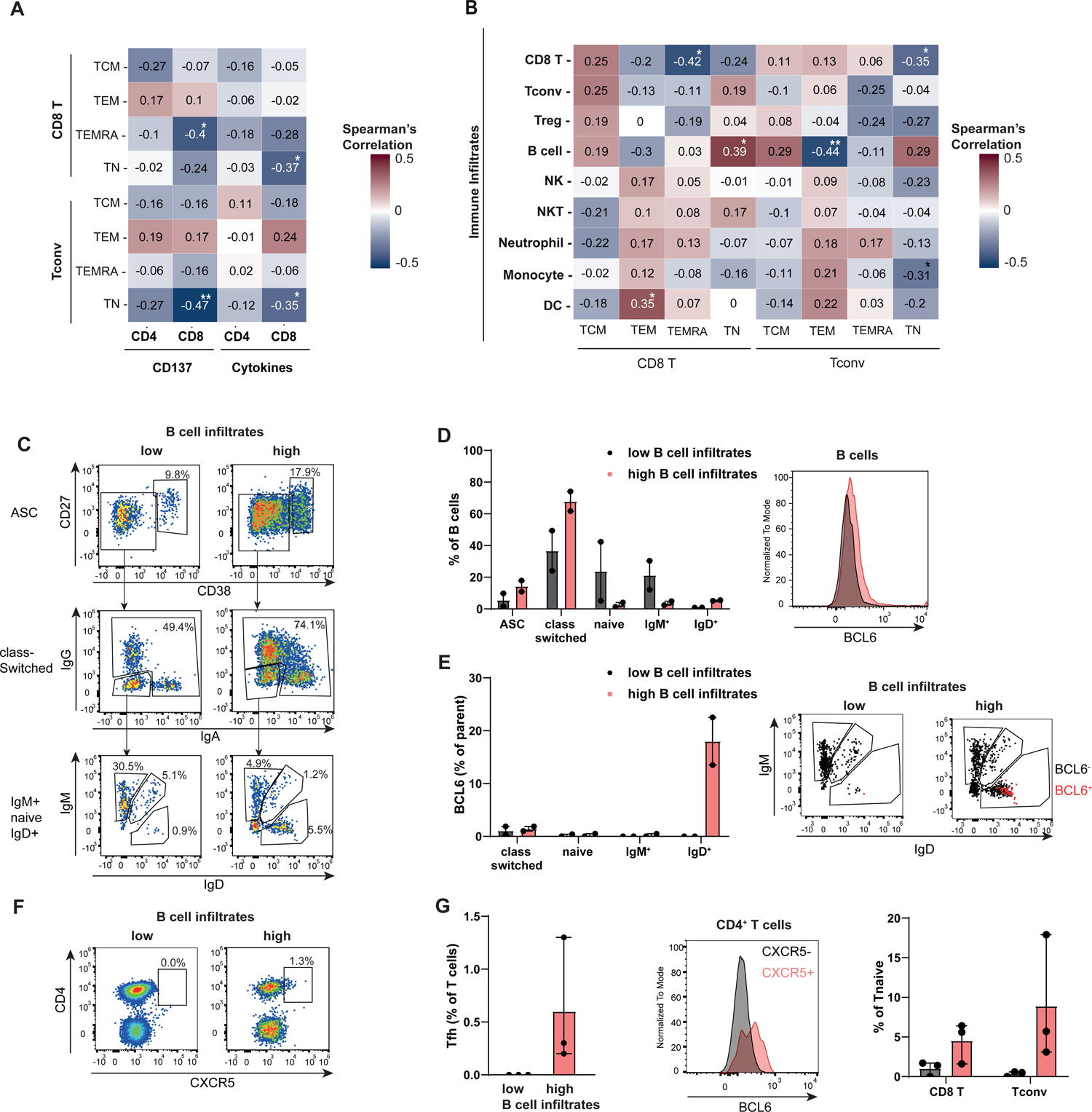
NSCLC lesions with high B cell infiltrates display TLS-specific immune cell composition. (A) Correlation heatmap of T cell differentiation subsets in the tumor digest with indicated anti-tumoral response of expanded TIL products upon 6h of co-culture with autologous tumor digest. (B) Correlation heatmap of different immune infiltrates with T cell differentiation subsets in the tumor digest. All correlation heatmaps report Spearman’s coefficient (r_s_) and *p<0.05, **p<0.005. (C) Representative gating on CD19^+^ B cells to identify CD27^+^ CD38^+^ antibody secreting cells (ASC), CD27^+^ class-switched B cells, IgM^+^ IgD^+^ naïve B cells, and IgM^+^ B, or IgD^+^ B cells. (D) Left panel: compiled data of patient #30 and #IV-14 with low B cell infiltrates, and patient #31 and #42 with high B cell infiltrates. Right panel: BCL6 expression of CD19^+^ B cells derived from NSCLC lesions with low (black) or high (red) B cell infiltration. (E) Left panel: Percentage of BCL6 expression on indicated B cell subsets. Right panel: Representative flow cytometry analysis of BCL6^+^ expression. (F) Representative Gating of CXCR5^+^ on CD4^+^T cells. (G) Left panel: percentage of BCL6 expression on Tfh from patient #30, #IV-14 and #IV-16 (low B cell infiltration) and patient #31, #42 and #IV-15 (high B cell infiltration). Middle panel: representative BCL6 staining of tumor-infiltrating Tfh. Right panel: percentage of naïve T cells present in indicated tumor lesions. Bar graphs depict mean with range.

Interestingly, when we correlated the immune infiltrates in the TME with the T cell differentiation status, we observed a positive correlation of B cell infiltrates with the presence of naïve CD8^+^ and CD4^+^ T cells, at the expense of Tem cells (figure 4B, supplemental figure 4C). Combined with the observation that B cell infiltrates positively correlated with CD4^+^ T cell infiltrates (figure 2A), yet negatively correlated with the presence of tumor-reactive T cells in the expanded TIL product (figure 2E), this finding let us to hypothesize that increased B cell infiltration might recruit naïve bystander T cells, which is indicative for TLS^52–54^.

To test this hypothesis, we characterized the B cell and CD4^+^ T cell tumor infiltrates in more detail. Of tumor digest we had available, we selected 2 patients with a low and 2 patients with a high percentage of B cell infiltrates. Intriguingly, tumors with high B cell infiltrates contained higher percentages of class-switched IgG^+^ and IgA^+^ expressing B cells, and very low percentages of naïve IgM^+^IgD^+^ B cells (figure 4C,D). Tumors with high B cell infiltrates also contained more antibody secreting cells (ASC), which is indicative of the presence of follicle formation and plasma cells^54^ (figure 4D). IgD^+^ B cells were almost exclusively present in tumors with high B cell infiltration (figure 4D).

To further define the type of B cell infiltrates, we measured BCL6 expression, which is indicative for germinal center B cells^55^. BCL6 expression on B cells was slightly increased in tumors with high B cell infiltrates (figure 4D, right panel). Notably, the BCL6 expression almost exclusively originated from IgD^+^ B cells (figure 4E), which entail mantle zone B cells^55^.

Concomitant with this finding, tumor lesions with high B cell infiltrates also contained CXCR5^+^ follicular helper T cells (Tfh, n=3) (figure 4F,G), with similar percentages as previously reported ^56,57^. Moreover, Tfh cells were undetectable in poorly B-cell infiltrated tumors (figure 4F,G). Importantly, BCL6 expression was exclusively detected in CXCR5^+^ CD4^+^ T cells, but not on CXCR5^-^ conventional CD4^+^ T cells (figure 4G), further consolidating that CXCR5^+^ CD4^+^ T cells are bona fide Tfh cells^58^. Lastly, in line with the correlation plots (figure 4B), we found more naïve CD4^+^ and CD8^+^ T cells in the tumors with high B cell infiltration (figure 4G). Thus, we conclude that high B cell infiltrates in NSCLC tumors indicate the presence of Tertiary lymphoid structure-like infiltrates, including mantle zone B cells, Tfh cells and naïve CD4^+^ and CD8^+^ T cells. This, in turn, negatively correlates with the tumor reactivity of expanded TIL products.

## Discussion

The immune cell composition of the tumor microenvironment is a strong indicator for survival of cancer patients^23–25^, and it can influence the success rate of immunotherapies^59–62^. Here, we show that the tumor-reactivity of expanded TIL products from NSCLC tumors negatively correlates with high B cell infiltration in the TME. When high percentages of B cells are measured, BCL6-expressing IgD^+^ mantle zone B cells and BCL6^+^ CXCR5^+^ follicular T helper cells are detected, which are both indicative of the presence of tertiary lymphoid structures^54,55,58^. In line with these findings, we detected higher percentages of naïve T cells in tumor digests with high B cell infiltrates.

Previous reports showed that the presence TLS structures- and thus high B cell infiltrates- are positive indicators for good response rates to checkpoint inhibitors^23,25^. Therefore, it may seem counterintuitive that high B cell infiltrates in the tumor digest negatively correlate with tumor-reactivity of expanded TIL products. We postulate that this divergence stems from the different starting point of these two therapeutic approaches. The anti-tumoral effect of checkpoint inhibitors depends not only on re-activation of tumor-specific T cells infiltrating the tumor^44,46,63,64^, but also on the activation of newly generated tumor-specific T cell responses. The latter occurs in draining lymph nodes and in TLS^65^, ensuring a continuous recruitment of novel T cell clones during the treatment. The therapeutic success of checkpoint inhibition thus relies on dynamic changes of immune cell infiltrates and activity in the TME. In contrast, the generation of tumor-reactive TIL products from tumor lesions depends entirely on the status quo of T cell infiltrates at the time of tumor excision. To obtain tumor-reactive TIL products, the tumor lesion must already contain tumor-reactive TILs, which can be expanded and reset to potent effector T cells during the 4 weeks expansion protocol of the TIL cultures. The success of TIL therapy thus depends on available T cell clones within the tumor lesions. It is thus tempting to speculate that whereas TLS support the activity of checkpoint inhibitors in the patients, a high prevalence of TLS-associated immune infiltrates, which increases the presence of naïve bystander T cells that still need to be primed^52,54^, rather hampers the effective outgrowth of tumor-reactive T cell during expansion. Combined, our findings strongly suggest that the parameters defining the treatment success of checkpoint inhibitors versus TIL therapy differ. In line with this hypothesis, recent studies showed that patients that were refractory to checkpoint inhibitors can still benefit from TIL therapy^7,15^.

We also observed other correlations of immune infiltrates with the tumor reactivity of TIL products. For instance, the presence of neutrophils positively correlated with TIL functionality. Interestingly, this positive correlation primarily originated from neutrophils that lacked the expression of PD-L1 or HLA-DR. PD-L1 expression on neutrophils has been shown to block T cell activation and function^66^ and correlates with poor outcome of hepatocellular carcinoma and gastric cancer patients^67,68^. Also in NSCLC, neutrophils are present in high numbers in the tumor lesions, and define the treatment outcome to immunotherapy^23,41,69^, as neutrophils are considered a negative prognostic factor for NSCLC patients treated with the checkpoint inhibitors^23,41,70,71^. Interestingly, we show here that high percentages of neutrophil infiltrates rather positively correlate with the tumor-reactivity in TIL products. This, however, only holds true when neutrophils express low levels of PD-L1 and HLA-DR. It would be interesting to further decipher the nature and heterogeneity of neutrophils within the tumor lesions. However, because neutrophils can only be phenotyped and further studied from freshly digested tumor lesions^72^, we could not follow up on this observation. Nevertheless, measuring the overall activation status of neutrophils may provide clues to define under which circumstances neutrophils are immunosuppressive.

In conclusion, we show here that the efficacy of generating tumor-reactive TIL products correlates with the immune cell composition of the obtained tumor lesion. The role of immune composition substantially differs from what is required for patients to respond to checkpoint inhibition. Therefore, we propose that defining the composition of immune infiltrates may help define whether a patient is more likely to respond to checkpoint inhibition, or to TIL therapy.

## Materials and Methods

### Tumor sample collection

Tumor samples were collected between April 2016 and November 2020. From early-stage patients, tissue from tumor lesions and healthy distal lung tissue was obtained (n=30). From late-stage patients, tissue from metastatic tumor lesions were collected (n=20). All tumor and tissue samples were measured in length at the pathology department. Four early-stage patients were excluded from the downstream analysis due to technical errors, resulting in a cohort of 26 early-stage NSCLC patients and 20 late-stage NSCLC patients. 11 early-stage and 18 late-stage patients had been included in previous reports^12,14^ (table 1). Sex was not considered as a biological variable. The study was performed according to the Declaration of Helsinki (seventh revision, 2013), with consent of the Institutional Review Board of the Netherlands Cancer Institute/Antoni van Leeuwenhoek Hospital (NKI-AvL), Amsterdam, the Netherlands. Tumor and lung tissue was obtained and processed within 4 hours after surgery.

### Tissue digestion

All samples were weighed prior to digestion. Tissue digestion was performed as previously described^12,14^. Briefly, freshly isolated tumor or lung tissue was finely chopped and enzymatically digested for 45 min at 37°C with RPMI (Gibco) containing 30 IU/ml collagenase IV (Worthington), 12,5 μg/ml DNAse (Roche), and 1% FBS (Bodego, Bodinco BV) to obtain a single cell suspension. Live and dead cells were manually counted with tryphan blue solution (Sigma) on a hemocytometer. Approximately 1×10^6^ cells were used per antibody staining mix for flow cytometry analysis. The remaining digest was used for TIL expansion cultures, or cryo-preserved until further use.

### TIL expansion

1-3×10^6^ cells were cultured in 24-well plates at 37°C and 5% CO2 in 20/80 T-cell mixed media (Miltenyi) containing 5% human serum (HS) (Sanquin), 5% FBS, 50 μg/ml gentamycin, 1.25 μg/ml fungizone, and 6000 IU human recombinant (hr) IL-2 (Proleukin, Novartis) for 10-13 days (pre-Rapid Expansion Phase; pre-REP). Cells were cultured for another 10-13 days (REP) with irradiated PBMCs pooled from 15 healthy blood donors, 30 ng/ml anti-CD3 antibody (OKT-3) (Miltenyi Biotec) and 3000 IU/ml hrIL-2 as previously reported^12^. Expanded TILs were used immediately for analysis, or cryo-preserved in RPMI medium 1640 containing 10% Dimethyl Sulfoxide (DMSO) (Corning) and 40% FBS until further use.

### T cell activation

Expanded TILs (REP) were counted with tryphan blue staining. To allow for detection of CD4 and CD8 expression after activation, cells were pre-stained with anti-CD4 and anti-CD8 for 30 min at 4°C in MACS buffer (PBS supplemented with 2% FBS and 2mM EDTA) and anti-CD107α was added at the start of the co-culture. Cells were washed once with MACS buffer and taken up in fresh 20/80 T cell mixed media. 1×10^5^ TILs were co-cultured with 2×10^5^ autologous tumor digest for 6-7 hours at 37°C. After 1h, 1x Brefeldin A (Invitrogen) and Monensin (Invitrogen) was added. As controls, expanded TILs were stimulated with 10 ng/ml phorbol myristate acetate (PMA) and 1 μg/ml ionomycin for 6-7h, or cultured with T cell mixed medium alone.

### Flow cytometry acquisition and analysis

Staining protocols for freshly digested tumor material or expanded TILs was previously described ^12,14^. For detailed information about antibodies, see table 2. Cells were washed twice with MACS buffer before staining. Live/dead fixable near-IR APC-Cy7 dye (Invitrogen) was included in surface staining mix for dead cell exclusion. Freshly digested single cell suspensions from tumor and lung tissue were stained with CD1a, anti-CD3, anti-CD11b, anti-CD11c, anti-CD14, anti-CD15, anti-CD16, anti-CD19, anti-CD20, anti-CD33, anti-CD45, anti-CD274, and anti-HLA-DR. For T cell differentiation analysis, single cell suspensions from digested tissue were stained with anti-CD3, anti-CD4, anti-CD8, anti-CD25, anti-CD27, anti-CD45RA, anti-CD127. Intracellular staining with anti-CD68 (freshly digested tumor lesions) and anti-Foxp3 (T cell differentiation) was performed after fixation for 30 min with Perm/Fix Foxp3 staining kit (ebioscience), according to the manufacturer’s protocol.

After T cell activation, cells were washed twice with MACS buffer and stained with anti-CD3 and anti-CD279. Live/dead fixable near-IR APC-Cy7 dye (Invitrogen) was included for dead cell exclusion. Cells were washed twice with MACS buffer and fixed for 30 min with Perm/Fix Foxp3 staining kit (according to the manufacturer’s protocol) and stained with anti-CD137, anti-CD154, anti-IL2, anti-IFNγ, and anti-TNF for 30 min at 4°C.

Cells were washed once with 1x Perm/Wash (ebioscience), taken up in MACS buffer and passed through a 70 µm single-cell filter prior to flow cytometric analysis on Symphony A5 (BD Biosciences). To ensure reproducibility, flow cytometry settings were defined for each patient with single antibody staining, and a standardized PBMC sample pooled from 4 healthy donors that was cryopreserved before the start of the study and used throughout.

Phenotypic analysis of tumor-infiltrating B cells and CD4^+^ T cells was performed on cryopreserved tumor digest patients with low B cell infiltrates (#30, #IV-14, #IV-16), patients with high B cell infiltrates (#31, #42, #IV-15). Defrosted cells were incubated for 1 hour at 37°C in RPMI at 2×10^6^ cells/ml containing 5% HS, 5% FBS, 50 μg/ml gentamycin, 1.25 μg/ml fungizone and 50 IU hrIL-2 (culture medium; CM), and then stained in MACS buffer for 30 min at 4°C with anti-CD3, anti-CD4, anti-CD11c, anti-CD19, anti-CD21, anti-CD24, anti-CD27, anti-CD38, anti-CD138, anti-IgA, anti-IgD, anti-IgG and anti-IgM, see detailed information in table 2. Live/dead fixable near-IR APC-Cy7 dye (Invitrogen) was included for dead cell exclusion. Cells were washed twice with MACS buffer and fixed for 30 min with Perm/Fix Foxp3 staining kit and stained with anti-CXCR5 and anti-BCL6^28^ for 30 min at 4°C, according to manufacturer’s protocol. Cells were washed once with Perm/Wash, taken up in MACS buffer and passed through a 70 µm single-cell filter prior to flow cytometry analysis on ID7000 Spectral Cell Analyzer (Sony Biotechnologies). Data analysis was performed with FlowJo Star 10.7.1 (BD).

### Gating in R

For flow cytometry measurements of T cell subsets and T cell differentiation status from tumors that had not been reported previously (table 1), and PD-L1 and HLA-DR expression on immune infiltrates were analyzed in R (version 4.1.1)^29^. FCS files were exported from the Symphony A5 (BD Biosciences) flow cytometer and imported into the R environment using flowCore package (version 2.4), and then cleaned with flowAI package (version 1.22.0)^30,31^. Cleaned FCS files were compensated using CytoExploreR package (version 1.1.0)^32^. Manual and automated gates with OpenCyto package (version 2.4)^33^ were used to gate for different T cell subsets and T cell differentiation states (supplemental figure 2A and 4A), as well as for PD-L1 and HLA-DR expression on immune infiltrates (supplemental figure 3A). The resulting gates were visualized with ggcyto package (version 1.20.0)^34^.

### Data processing

To define the overall lymphoid and myeloid infiltrates from tumor lesions and lung tissue, FCS files from FACSDiva were exported, compensated and pre-gated in FlowJo Star v10.7.1 to remove debris, doublets, dead cells, and to enrich for CD45^+^ live cells. Pre-gated FCS files were exported from FlowJo and pre-processed in a multistep analysis with an in-house developed R pipeline (https://github.com/NandhiniKanagasabesan/NSCLC_Project). First, FCS files were imported into the R environment, and expression values of selected markers were extracted from each FCS file with flowCore package (version 2.4) (supplemental figure 1B)^35^. Batch correction for all patient combined was performed as follow: 1) The marker expression values were transformed using hyperbolic arcsine transformation, which linearizes near-zero values and logarithmically transforms higher positive and negative values with rrscale package (version 1)^30,36^ and 2) Transformed data were then median-centered and scaled^30^.

### Clustering in CytoTree

Pre-processed FCS files were clustered and visualized with CytoTree package in R (version 1.3)^37^. For UMAPs, the expression values from each pre-processed FCS file were merged using the ceil method. Pre-processed FCS file with more than 50k cells, with sub-sampled at random 50k cells (without replacement); otherwise, all data was used and aggregated to allow for downstream application. The expression matrix and metadata (patient ID) were combined to build a CYT object, on which unsupervised clustering was performed with self-organizing maps (SOM), resulting in 25 clusters. Clusters were processed by dimensionality reduction (UMAP). Tree-shaped trajectories were built using minimum spanning tree (MST) approach. With heatmap visualization and tree plot, clusters were assigned to meta clusters (immune cell populations) that were identified by comparing the expression of markers (supplemental figure 1B). Clusters of immune cell populations were visualized on a 2-dimensional UMAP plot.

### Statistical Analysis and Data visualization

The percentage of live immune infiltration was calculated with CytoTree package in R. The difference between lung and early-stage lesions was calculated with paired Student t-test, and between early stage and late-stage lesions with unpaired Student t-test. Correlation of immune infiltrates with immune infiltrates, TIL expansion and with tumor reactivity was calculated using Spearman’s Rank correlation with Hmisc R package (version 4.7.1)^38^. The correlation was considered statistically significant when p-value < 0.05. Plots, correlation heatmaps and raincloud plots were generated with GraphPad Prism (Dotmatics, version 9.1.1), Cytotree (version 1.3), ggplot2 (version 3.4.2)^37,39^, and ggdist (version 3.3)^40^ packages in R.

## Declarations

### Authors’ contributions

MvL, RdG, SC and MW designed the project. AG, MvL, RdG and SC performed the practical work, SC and NK performed data analysis. SC and MW wrote the manuscript. All authors provided input on the manuscript. MW supervised the study.

## Acknowledgements

We like to thank the medical assistance staff from the NKI-AvL and the flow cytometry facility from Sanquin research for technical help, and all patients contributing to this research project. We also thank N. Verstegen, L. Kuijper, L. Fernandez Blanco, L. Kummer and A. ten Brinke (Sanquin Blood supply) for support with the B cell analyses. We thank K. Bresser for support on the bioinformatic data analysis.

