## Supplemental figure 1 for "Tertiary lymphoid structure-related immune infiltrates in NSCLC tumor lesions correlate with low tumor-reactivity of TIL products"

A

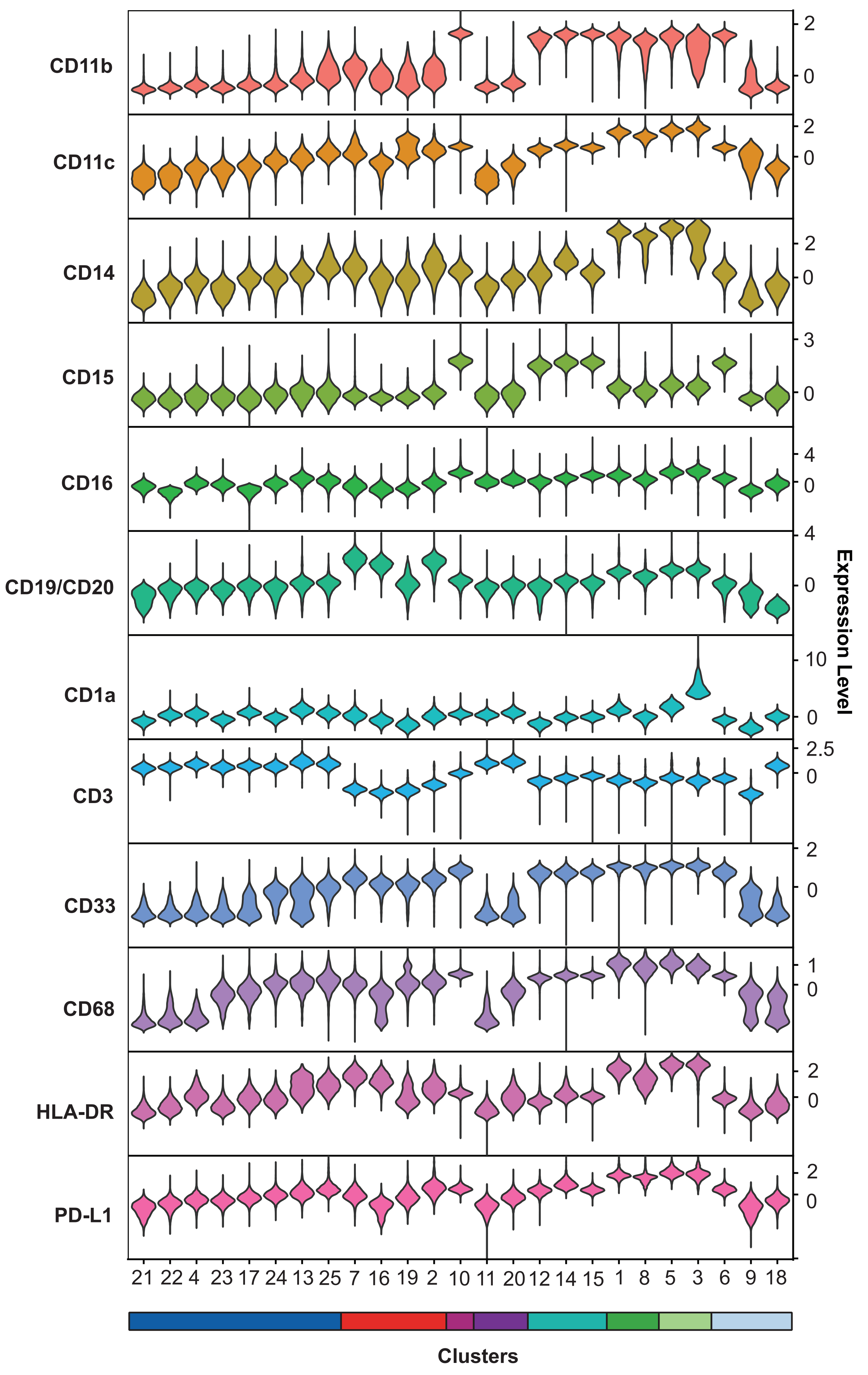


B

| Immune cell populations | Markers: Inclusion | Markers: Exclusion |
|---|---|---|
| T cells | CD3 | CD11b, CD14, CD15, CD16, CD19/CD20 |
| B cells | CD19/CD20 | CD3, CD11b, CD14, CD15, CD16 |
| NK cells | CD16 | CD3, CD11b, CD14, CD15, CD19/CD20 |
| NKT cells | CD3, CD16 | CD11b, CD14, CD15, CD19/CD20 |
| Neutrophils | CD11b, CD15 | CD3, CD14, CD19/CD20 |
| Monocytes | CD11b, CD14, CD33, CD68, HLA-DR | CD3, CD15, CD19/CD20 |
| Dendritic cells | CD1a, CD11c, PD-L1, HLA-DR | CD3, CD14, CD15, CD19/CD20 |

C

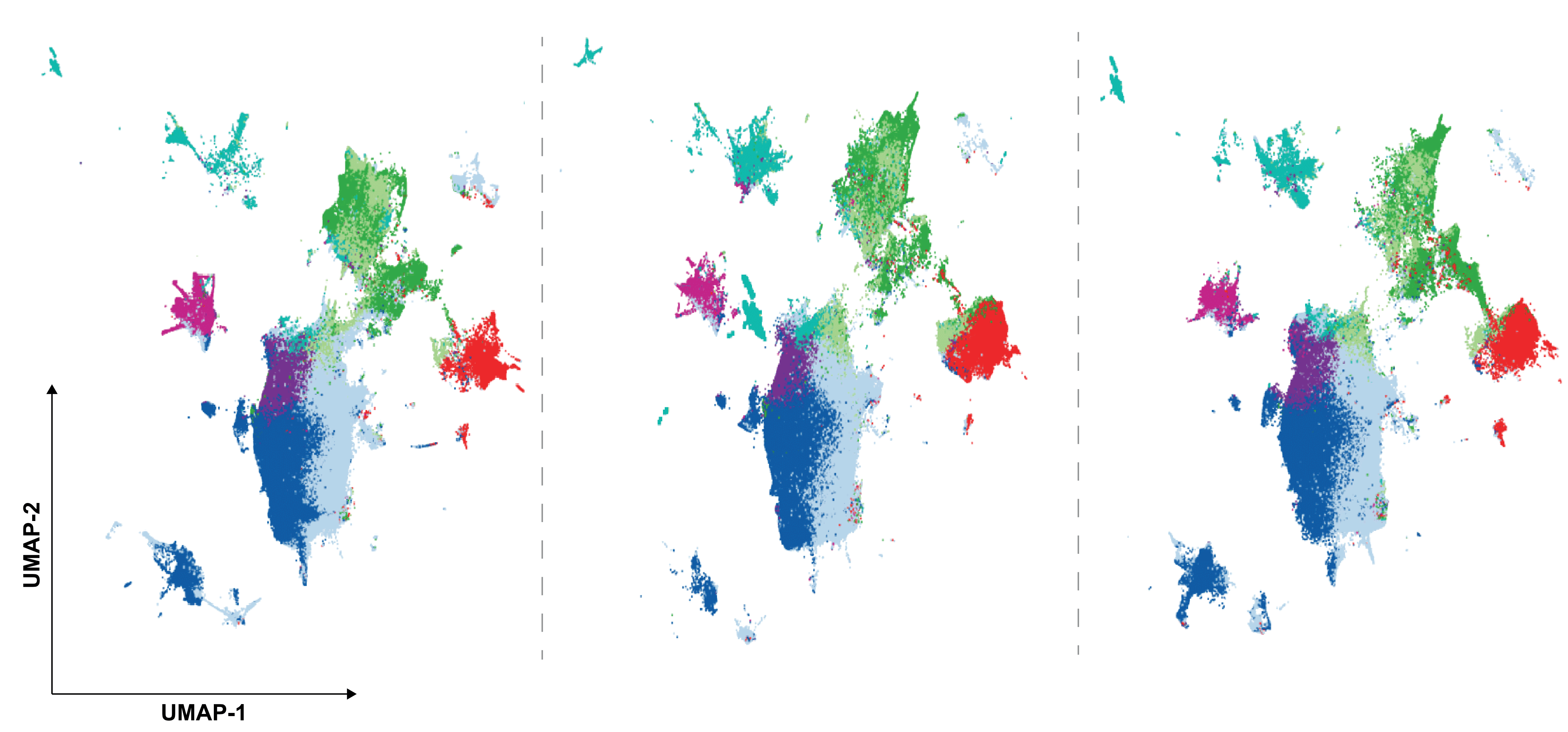


**Supplementary Figure 1. Lymphocyte and myeloid cell populations in healthy adjacent lung tissue and NSCLC tumor lesions.**
(**A**) Violin plots indicating the expression distribution of markers in each cluster. The colored bar below depicts the clusters assigned to their respective immune cell populations by comparing the marker expression summarized in (**B**), and light blue indicates undefined CD45$^{+}$ immune infiltrates. (**C**) UMAP projections of different CD45$^{+}$ immune cell subsets found in the healthy lung digest, early-stage and late-stage NSCLC tumor digest. Each dot corresponds to a single cell, colored in accordance to cell type.
