## Supplemental figure 2 for "Tertiary lymphoid structure-related immune infiltrates in NSCLC tumor lesions correlate with low tumor-reactivity of TIL products"

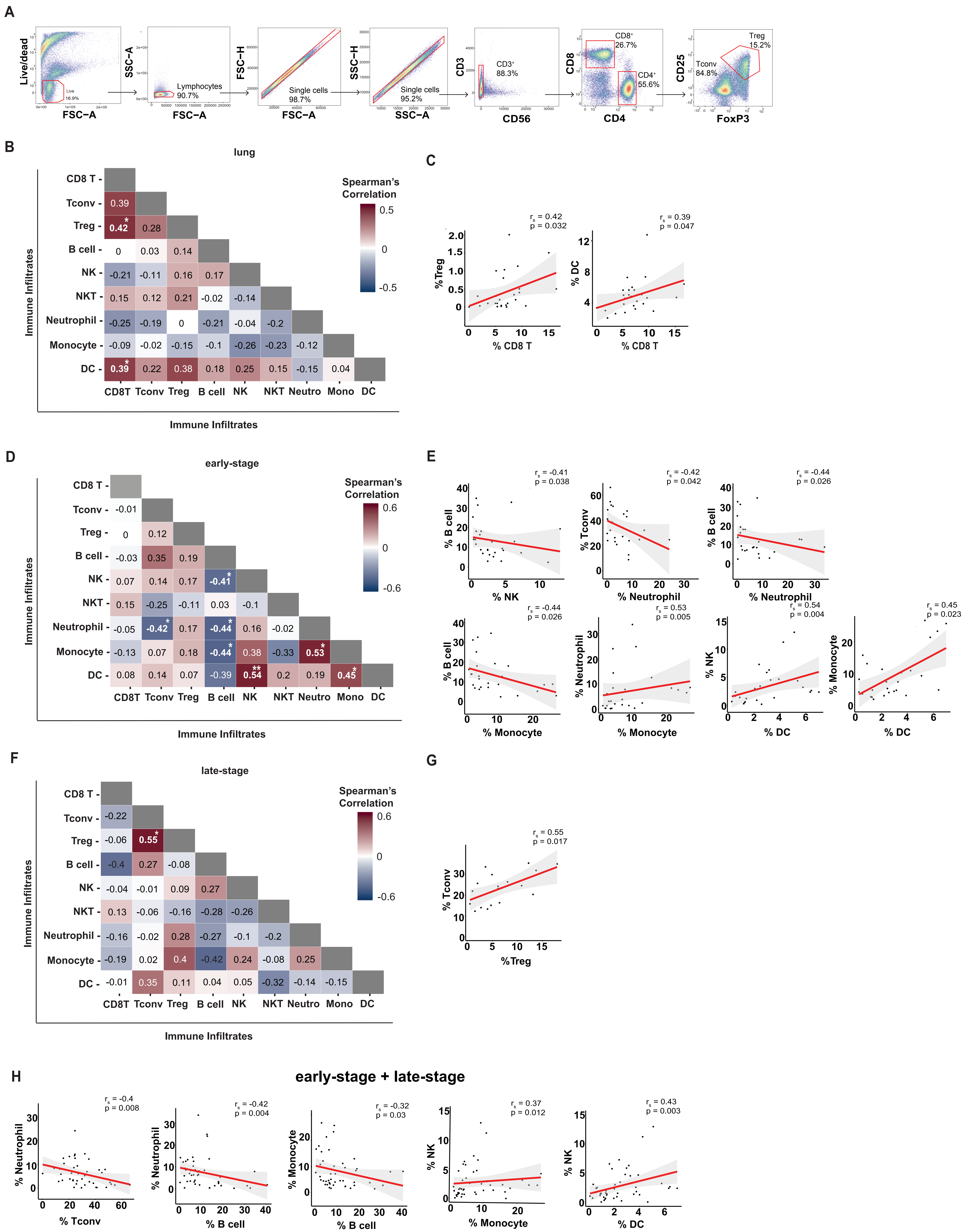

**Supplementary Figure 2. Correlation between Immune cell subsets in lung tissue and in early and late stage NSCLC immune infiltrates.**
(**A**) Example of gating strategy for defining CD3+ T cell subsets (patient #31) for correlation analysis in Figure 2. Correlation heatmap between different immune cell infiltrates of (**B**) lung tissue (n=26), (**D**) early-stage NSCLC (n=26) and (**F**) late-stage NSCLC lesions (n=20). Significant correlations are shown as scatterplots in **C**, **E**, **G**. (**H**) Scatterplots show significant correlations of Figure 2A. All correlation heatmaps report Spearman's coefficient ($r_s$) and *p<0.05, **p<0.005. All scatterplots show the Spearman's coefficient ($r_s$) and p-value. The red line indicates the regression line of the correlation, and the grey area represents the confidence interval.
