## Supplemental figure 3 for "Tertiary lymphoid structure-related immune infiltrates in NSCLC tumor lesions correlate with low tumor-reactivity of TIL products"

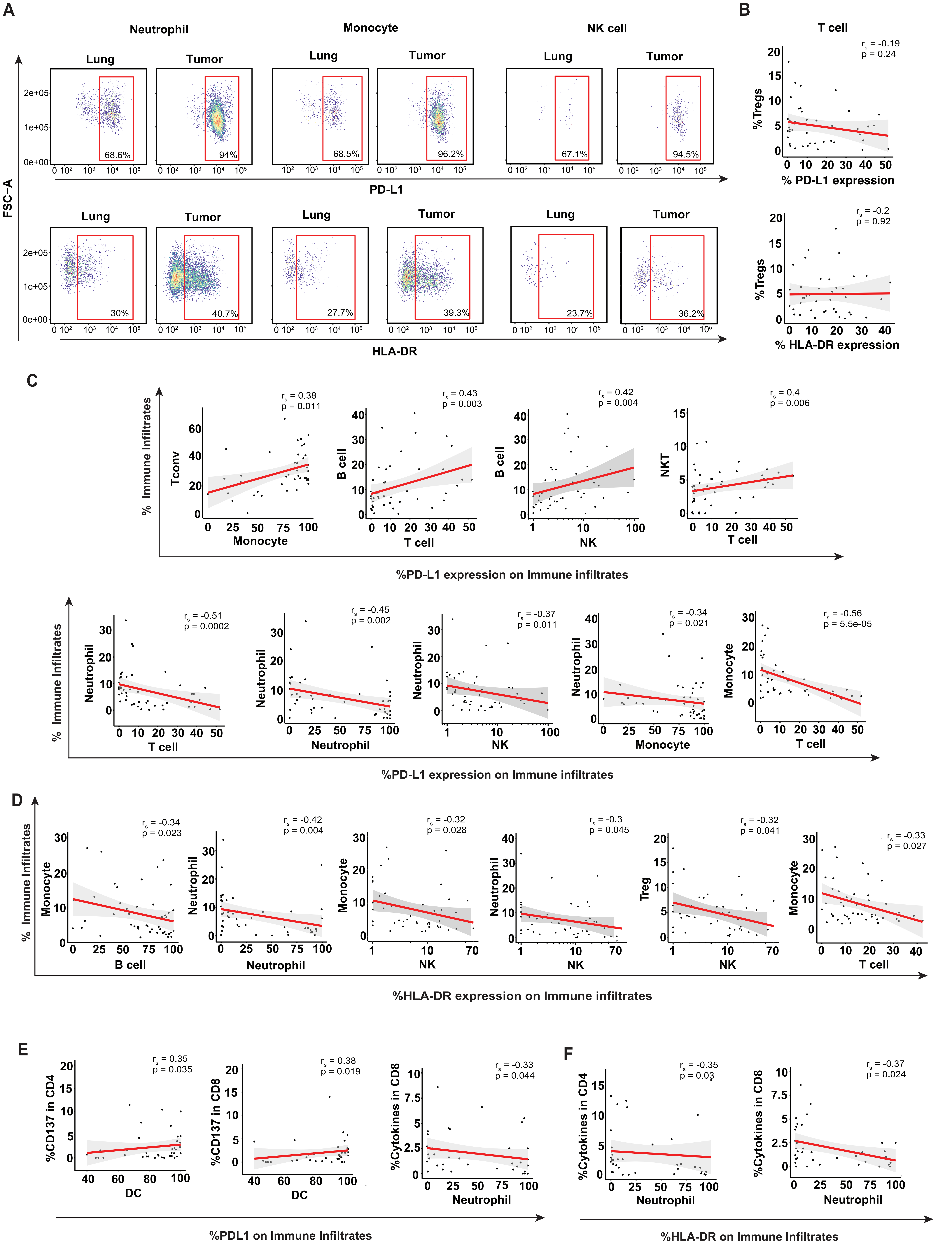


**Supplementary Figure 3. Correlation of PD-L1 and HLA-DR expression on immune infiltrates with overall infiltrates and with functionality of TIL products.** (**A**) Gating strategy for PD-L1 (top-row) and HLA-DR (bottom-row) expression on neutrophils, monocytes, and NK cells (patient #51). (**B**) Scatterplots of the percentage of PD-L1 expression (top) and HLA-DR expression (bottom) on T cells with the percentage of Tregs infiltrating NSCLC lesion. (**C, D**) Scatterplots show the significant correlations of Figure 3C, D; (**C**) the percentage of PD-L1 expression or (**D**) the percentage of HLA-DR expression on indicated immune infiltrates (x-axis) with the percentage of immune infiltrate in NSCLC tumor lesions (y-axis). (**E, F**) Scatterplots show the significant correlations of Figure 3F; (**E**) the percentage of PD-L1 expression or (**F**) the percentage of HLA-DR expression on indicated immune infiltrates (x-axis) with the percentage of T cell functionality of expanded TIL products (y-axis). All scatterplots report Spearman's coefficient ($r_s$) and p-value. Each dot represents a patient. The red line indicates the regression line of the correlation, and the grey area represents the confidence interval.
