## Supplemental figure 4 for "Tertiary lymphoid structure-related immune infiltrates in NSCLC tumor lesions correlate with low tumor-reactivity of TIL products"

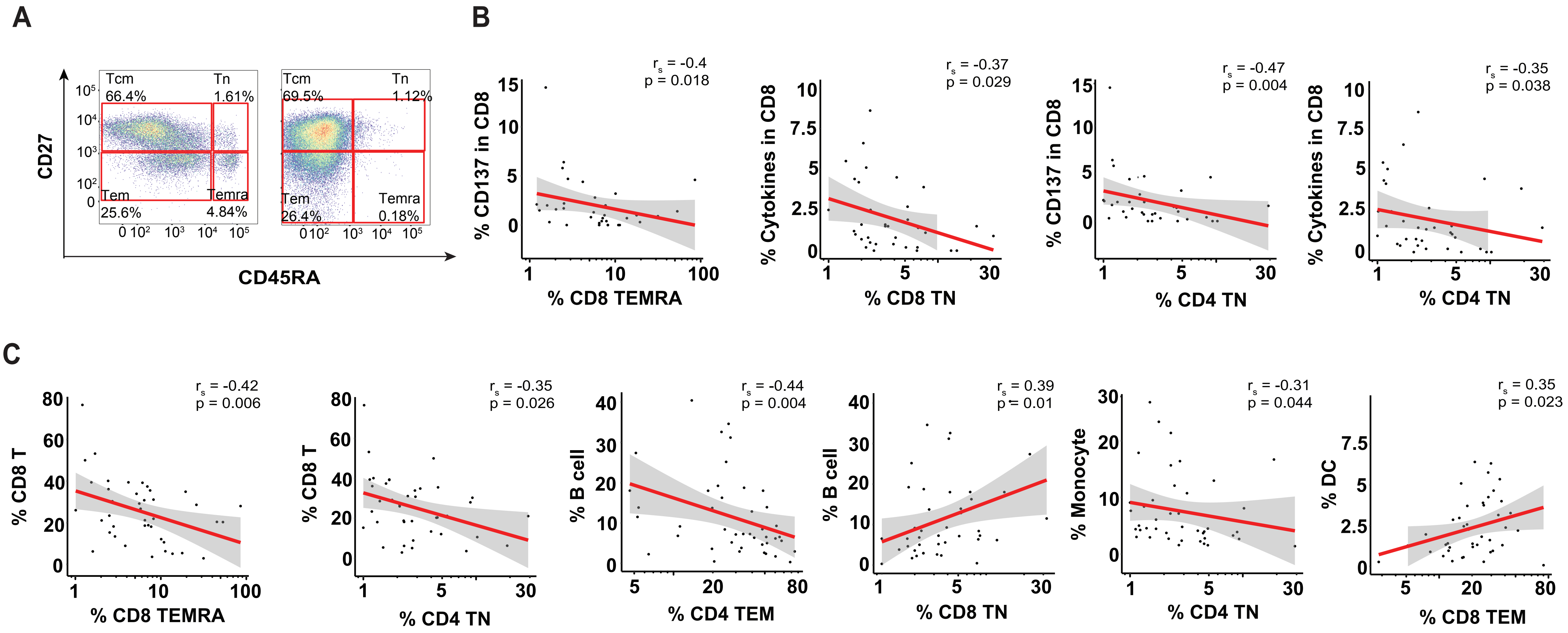


**Supplementary Figure 4. Correlation of T cell differentiation status with immune infiltrates in NSCLC tumor lesions and with T cell functionality of TIL products**
(**A**) Gating strategy for T cell differentiation subsets (Tcm, Tem, Temra, and Tn) on CD8+ and conventional CD4+ T cells (patient #31). (**B**) Scatterplots show the significant correlations of Figure 4A; the percentage of T cell differentiation subsets in NSCLC tumor lesions (x-axis) with the percentage of T cell functionality of expanded TIL products (y-axis). (**C**) Scatterplots show significant correlations of Figure 4B; the percentage of T cell differentiation subsets in NSCLC tumor lesions (x-axis) with the percentage of different immune infiltrate in NSCLC tumor lesions (y-axis). All scatterplots report Spearman's coefficient ($r_s$) and p-value. Each dot represents a patient. The red line indicates the regression line of the correlation, and the grey area represents the confidence interval.
